# Quantitative Analysis of *Meloidogyne incognita* Population Density Using Real-Time PCR and Its Correlation with Root-Knot Disease Index in Tabacco (*Nicotiana tabacum*)

**DOI:** 10.1101/2025.08.27.672540

**Authors:** Zejun Cheng, Jingwen Chai, Haoshuai Pu, Zhe Zhao, Xiaoxin Duan, Wei Zheng, Jianqiang Xu, Pu Miao, Wenbang Hou

## Abstract

Tobacco root-knot disease represents a significant threat to tobacco production, particularly in the western Henan region, where *Meloidogyne incognita* is the predominant species. This study collected samples of *M. incognita* and soil from Luoyang, Henan, and designed specific primers MiF and MiR based on the amplified 735 bp sequence of the ITS1-5.8S-ITS2 region. These primers exhibit mismatches with related species, including *M. javanica, M. minor, M. hapla*, and *M. arenaria*, showing 1, 6, 10, and 10 base differences in the forward direction, respectively, 3, 10, and 9 base mismatches in the reverse direction. Although the primers were used to detect corn-wheat soil samples, no amplification was observed. Additionally a real-time quantitative PCR curve for *M. incognita* in soil was constructed, revealing a negative correlation between the Ct value (Y) and the log-transformed number of nematodes (x) per 20 g of dry soil, represented by the equation y = -0.9757x + 35.565; (R^2^ = 0.9999, *P* < 0.01). According to the decomposition efficiency of nematode DNA in soil, the results showed that nematode DNA degrades rapidly in soil, with a degradation rate of approximately 87.3% at 3 days and 99.97% at 14 days. Furthermore, significant differences were observed in the real-time PCR detection efficiency among various nematode forms: the Ct value of J1 was significantly higher than that of J2, while abnormal eggs (empty eggs or internal bubble eggs) exhibited the highest Ct value. The proportion of abnormal eggs in the soil before planting was significantly higher at 64.08% compared to only 15.3% at harvest, indicating that the activity of nematodes in the soil is significantly reduced after harvesting in October and planting in March of the following year. A survey of 126 tobacco plants indicated a significant positive correlation between the root-knot index (RKI) and root-knot nematode density (r = 0.80, p < 0.01). The study identified a minimum disease threshold of 234 individuals / 20 g soil at harvest and revealed a nonlinear relationship between disease severity and nematode density. Specifically, a weak correlation was observed at low density (Low RKI: 0-1; nematode density<2000 individuals/20 g soil, r = 0.49, P<0.05), while a significant correlation was noted at moderate density (High RKI: nematode density>2000 individuals/20 g soil, r = 0.69, P<0.05). This study showed that as the RKI increases, the rate of increase in nematode density in the soil diminishes. These findings provide valuable insights for the development of effective scientific strategies for nematode control.

## 1. Introduction

Tobacco (*Nicotiana tabacum* L.) is the primary raw material for cigarettes and constitutes a significant economic crop in China [1]. As the largest producer and consumer of tobacco globally, China accounts for over one-third of the world’s tobacco production [2]. Major pests and diseases affecting tobacco include the tobacco mosaic virus (TMV), black shank (*Phytophthora nicotianae*), bacterial wilt (*Ralstonia solanacearum*), and root-knot nematode [3]. Notably, root-knot nematodes cause wounds on the epidermal cells of tobacco roots, which increase the susceptibility of tobacco plants to soil-borne pathogens. Therefore, controlling tobacco root-knot nematodes is essential, as it may reduce the incidence of other soil-borne diseases[4].

*Meloidogyne* spp. are widely distributed plant pathogenic nematodes that significantly affect the growth and yield of various economically important crops, including tobacco, tomato, and cucumber [10]. By invading the root system of plants, these nematodes induce the formation of root galls, disrupting the normal structure and function of the roots [11]. This disruption leads to a decline in water and nutrient absorption, resulting in stunted growth, yellowing of leaves, reduced yields, and in severe cases, plant death [12]. Statistics indicate that agricultural losses caused by root-knot nematodes worldwide amount to billions of dollars [13]. Tobacco root-knot diseases typically reduce yields by 10% to 20%, however, in severe cases, losses can exceed 75%, with a higher incidence of disease reported in developing countries compared to developed ones [10]. In provinces such as Henan, Anhui, Sichuan, Guizhou, Zhejiang, and Yunnan in China, tobacco root-knot disease affects approximately 52,000 hm^2^ annually, accounting for about one-tenth of the total production area, leading to losses of around 9 million dollars [14]. Therefore, early detection and effective prevention and control of tobacco root-knot nematodes are critical issues that must be addressed in agricultural production [15].

Currently, there are no tobacco varieties exhibiting high resistance to root-knot nematodes, and the interactions between these pests and host plants are complex, posing significant challenges for resistance breeding [5]. Consequently, root-knot nematode disease has emerged as a significant constraint on tobacco production in various countries and regions [6]. Presently, the most effective strategy for managing tobacco root-knot nematode disease focuses on suppressing the reproduction and population density of the pathogenic nematodes [7]. For instance, Zhang et al., cloned the tobacco RKN resistance gene NtRk1, which is induced upon nematode infection and enhances resistance by regulating salicylic acid and jasmonic acid pathways. Overexpression of this gene confers protection in susceptible cultivars while RNA interference (RNAi) silencing increases susceptibility to *Meloidogyne incognita* [8]. Additionally, Li et al. identified 5,206 high-confidence long non-coding RNAs (lncRNAs) in tobacco, with 565 being differentially expressed during nematode infection [9].

Traditional nematode detection methods primarily rely on morphological identification and microscopic observation [16]. Morphological identification necessitates a comprehensive understanding of the morphological characteristics of nematodes, as even minor differences among species can lead to misidentification [17]. Moreover, microscopic observation is time-consuming and often does not meet the rapid detection requirements for large-scale samples.

Conventional quantitative methods are inadequate for detecting nematode eggs, J1 and inactive nematodes in soil, resulting in unreliable disease forecasts [18]. With the rapid advancements in molecular biology technology, real-time fluorescent quantitative PCR (Real-Time PCR) has become widely utilized in plant disease detection due to its high sensitivity, specificity, and quantitative capabilities [19]. This technology allows for the rapid and precise detection of nematode DNA through the design of specific primers and fluorescent probes, facilitating accurate quantification of nematode density [21]. Compared to traditional morphological methods, real-time PCR technology significantly reduces detection time and effectively distinguishes between different types of nematodes, thereby providing robust technical support for early warning and precision prevention and control of diseases [21-24].

While real-time PCR technology demonstrates a significant advantage in detecting nematodes, current research remains limited [25]. The root-knot disease index (RKI) serves as a crucial indicator of disease severity, typically assessed through parameters such as the number of root knots and the extent of root damage [26]. Clarifying the quantitative relationship between root-knot nematode density and the disease index not only enhances our understanding of the infection mechanism of root-knot nematodes but also provides a scientific foundation for early diagnosis and comprehensive disease management. For instance, developing a mathematical model that correlates nematode density with disease severity enables the prediction of disease progression and the formulation of targeted prevention strategies [27]. This approach can minimize pesticide use, reduce environmental pollution, and enhance the sustainability of agricultural production [28].

The purpose of this study is to use real-time PCR technology for the quantitative detection of tobacco root-knot nematode density and to analyze the correlation between nematode density and the root disease index. This research will explore the manifestations of disease in the tobacco root system under varying nematode densities, thereby establishing a quantitative relationship between nematode density and the disease index. The findings will provide a theoretical foundation and technical support for early diagnosis, as well as comprehensive prevention and control of tobacco root diseases. Furthermore, the results of this research object not only to enhance the detection efficiency of tobacco root-knot nematodes but also to serve as a reference for the prevention and control of similar issues in other crops.

## 2. Materials and methods

### 2.1 Soil and nematode collection

The soil and tobacco samples were collected from four tobacco planting field in Luoyang: Luoning Xiaojie (111.59°E, 34.45°N), Yiyang Gaocun (111.88°E, 34.55°N), Ruyang Baipo (111.70°E, 34.50°N), and Kaiyuan Farm (112.42°E, 34.60°N). Prior to tobacco cultivation in March 2022, soil samples were collected from fields affected by root-knot nematodes at a depth of 0-30 cm. Egg masses of root-knot nematode in the soil were extracted and counted before transplantation. The Cobb’s sieving and decanting method, along with the flotation method, were employed for the collection of egg masses from the soil [29]. A total of 100 g of soil samples were taken, to which an appropriate amount of distilled water was added. This mixture was then filtered through 850 µm and 250 µm sieves to remove larger particle impurities and collect the residues.

The residue was re-suspended in a sucrose solution (454 g/L, with a density of approximately 1.18 g/cm^3^). After mixing, the solution was subjected to centrifugation at 1207 x g for 5 min. Due to the lower density of the egg masses, they floated to the surface of the sucrose solution, while heavier impurities settled at the bottom of the tube. A straw was used to carefully extract the liquid, which was then transferred to a new centrifuge tube. The mixture was centrifuged again to remove the sucrose, resulting in the isolation of the egg mass sample. On October 7, 2022, and October 15, 2023, tobacco root samples were collected following the tobacco harvest. Using a five-point sampling method, samples were taken from the 0-30 cm soil layer within a 50 cm radius centered on the tobacco plants. The tobacco root egg masses were separated from the tobacco roots, the fibrous roots were cut, and the egg within the root egg masses were extracted under a microscope.

### 2.2 Extraction of M. incognita DNA and Design of Specific Primers

The nematodes used for the identification of the DNA sequence of root-knot nematodes were collected from infected roots of tobacco plants in four different tobacco planting fields in October 2021. The egg masses were extracted from the root knots, and a single egg was used as a sample for DNA extraction. Using tweezers, the egg mass was broken to release the eggs and juvenile second-stage nematodes (J2). The eggs were then incubated at room temperature (25°C) for 3 h. A pipette was used to extract a single egg or J2, following the single nematode DNA extraction method described by Wang et al. (2011). A single J2 sample was placed on a sterilized glass slide alongside a drop of water. A No. 3 insect needle was then used to make a vertical incision in the J2 sample. Subsequently, 10 µL of the nematode solution was aspirated into a 200 µL centrifuge tube, and 1.5 µL of 10 x PCR Buffer (Mg^2+^-free) was added. The mixture was subjected to liquid nitrogen for 1 min, followed by heating at 85°C for 2 min. Then, 1 µL of 1 mg/mL protease K was incorporated, and the solution was heated at 56°C for 15 min followed by an additional 10 min at 95 °C. This DNA extraction solution was subsequently diluted 10 times with sterile water to serve as a template for PCR amplification.

The amplification of the ITS1-5.8S-ITS2 region was conducted using the forward primer ITS-F (5’-ACA AGT ACC GTG GAA AGT TG-3’) and the reverse primer ITS-R (5’-TCG GAA GGA ACC TAC TA-3’). PCR amplification was performed with Hieff Canace Plus High-Fidelity DNA Polymerase (Yeasen, Shanghai) under the following conditions: 94°C for 2 min, (94°C for 1 min, 48°C for 1 min, 72°C for 1 min) x 35 cycles, 72°C for 5 min. The PCR products were purified using the DiaSpin column PCR product purification kit (Sangon Biotech, Shanghai) and subsequently sent to Sangon Biotech for sequencing. The ITS region sequence of *M. incognita* obtained in this study was compared with sequence in the NCBI database for identification. The target sequence was analyzed using Mega 11.0 software to identify variant regions, and NCBI Primer-BLAST was employed to design primers specific to the nematode (MiF and MiR).

### 2.3 Establishing a real-time PCR method for the quantification of tobacco root-knot nematode

The DNA solution extracted from a single J2 was diluted ten fold with sterile distilled water to serve as a template for Real-Time PCR. Real-Time PCR was conducted using the CFX96 (Bio-Rad, USA) for detection. The reaction mixture consisted of 10 µL, which included 5 µL of Hieff qPCR SYBR GREEN Master Mix (Yeasen, Shanghai), 0.4 µL of specific primers, and 2 µL of template DNA. The reaction conditions were as follows: 95°C for 30 s; (95°C for 5 s and 58°C for 30 s) x 40 cycles. Distilled water was used as a negative control. Additionally, the DNA solution from a single *M. incognita* was diluted to 10^-1, 10^-2, 10^-3, 10^-4, and 10^-5 for use as templates, and specific primers MiF and MiR were employed across three replicates for treatment.

Three soil samples were collected from corn-wheat crop rotation fields in Yiyang Gaocun, following the method described by Cheng et al. (2018) for extracting total biological DNA from the soil. 20 g of soil samples were dried at 60°C for 24 h and processed using a planetary ball mill (LANENDE, Shandong) at a speed of 450 rpm for 2 min [30]. Subsequently, 5 g of the ball milled soil samples were added to 50 mL centrifugal tubes, along with 10 mL of phosphate buffer (Na_2_HPO_4_ and NaH_2_PO_4_, 0.12 M, pH 8.0), with three replicates for each sample. The soil-buffer mixture was agitated using a shaker (Hongke Technology HQY-C, Jiangsu, China) at 200 rpm for 30 m. After an additional 10 min, the solution was purified using the DiaSpin column PCR product purification kit (Sangon Biotech, Shanghai). The DNA eluate was then diluted ten-fold to serve as a template for real-time PCR. All samples were prepared in triplicate.

The DNA solution extracted from the soil sample of the corn-wheat crop rotation was detected using the MiF and MiR primers; however, no specific amplification was observed (data not shown). Consequently, a quantitative curve for the root-knot nematode of tobacco was created using soil collected from the corn-wheat rotation field. The tobacco roots containing egg masses were cut into 1 cm segments with scissors, and subsequently homogenized in a blender for 30 seconds (Midea WBL2521H, Guangdong). The fragmented roots were placed in a gauze bag, and both J2 and adults root-knot nematodes were extracted using the Baermann funnel method for inoculation. The soil from the corn-wheat rotation field was dried at 60°C and then weighed to a mass of 20 g. Different quantities of *M. incognita* (J2 and adults) were artificially inoculated: 0, 30, 150, and 750 individuals. Following inoculation, the soil was dried again at 60°C for 6 h. Each inoculation amount was prepared in triplicate.

### 2.4 Rate of DNA degradation of M. incognita in soil

In October 2023, tobacco roots infected with root-knot nematode were collected post-harvest. The infected roots were homogenized using a blender (Midea WBL2521H, Guangdong) for 30 s, with this process repeated twice. The nematode suspension, comprising both J2 and adult stages was subsequently extracted using the Baermann funnel method. Following treatment of the *M. incognita* suspension in a water bath at 75°C for 10 minutes, approximately 2000 individuals of *M. incognita* were inoculated into 20 g of raw soil sourced from a corn-wheat rotation field using a pipette for precision. DNA extraction from the soil was performed at intervals of 0, 3, 7 and 14 days, employing the DNA extraction method outlined by Cheng et al. (2018)[]. The extracted DNA was then analyzed through real-time PCR detection, yielding Ct values with the MiF and MiR primers. The equivalent number of nematodes was calculated based on the quantitative curve of *M. incognita*.

### 2.5 Effect of the tobacco root-knot disease index (RKI) on nematode population density (NPD)

On October 7, 2022, and October 15, 2023, samples were collected from four tobacco-growing fields in Luoyang City, Henan Province, specifically at Luoning, Yiyang, Ruyang Bai, and Kaiyuan Farm, following the tobacco harvest. At each location, 38, 32, 30, and 26 tobacco plants were sampled for roots, resulting in a total of 126 plants. Soil samples were also collected from the rhizosphere area (15 cm in diameter with the root; 0-30 cm in depth) of each tobacco plant and stored in sterile sealed bags. The established real-time PCR method for detecting *M. incognita* was used to employed to assess the nematode population density (NPD) in the soil samples. The diseased tobacco roots were washed with water to eliminate any adhering soil, and 30 g of root samples were cut and placed on a white tray for root-knot quantification. The number of galls was observed and counted either with the naked eye or using a magnifying glass. Concurrently, the Root-Knot Disease Index (RKI) of each tobacco plant was evaluated. Based on the number of root egg masses and the degree of root damage, a 0-5 scale grading standard from Taylor (1978) was used to evaluate the RKI of each tobacco root, with specific grading criteria as follows [31]:

0 level: No root-knots present;

1 level: 0% ≤ number of swollen roots or root-knots due to nematode damage < 10%;

2 level: 10% ≤ number of swollen roots or root-knots due to nematode damage < 25%;

3 level: 25% ≤ number of swollen roots or root-knots due to nematode damage < 50%;

4 level: 50% ≤ number of swollen roots or root-knots due to nematode damage < 75%;

5 level: 75% ≤ number of swollen roots or root-knots due to nematode damage ≤ 100%.

The infected young roots develop spherical or irregular galls of varying sizes, which appear white or pale in color. In severe cases, the diseased roots turn brown and decay. Under conditions of high RKI level, the root system becomes severely deformed, exhibiting a beaded or claw-like morphology.

### 2.6 Statistical analysis

Statistical analyses were performed using SPSS version 26.0. A linear regression analysis was conducted to explore the relationship between the number of inoculated nematodes and the Ct values obtained from real-time PCR, with a correlation coefficient calculated (p < 0.01). Furthermore, the Pearson correlation coefficient was employed to assess the correlation between the root-knot disease index (RKI) and the nematode population density (NPD).

## 3. Result

### 3.1 Identification of M. incognita and Design of Specific Primers

The findings of this study indicate that the ITS region sequences of root-knot nematodes from tobacco in four planting areas of Luoyang, Henan Province, exhibit a high degree of consistency. Analysis using NCBI BLAST revealed a similarity of 99.57% with *M. incognita* from Fujian Province, China (accession number: OQ632600), with only three base pair mismatches in a 735 bp sequence. Additionally, a similarity of 99.28% was observed with the sequence designated as accession number MT209949, which contained five base pair mismatches. For further analysis, the sequence was compared with those of both closely related and distantly related species, as provided by the National Center for Biotechnology Information (NCBI) (Figure 1).

**Figure 1.**
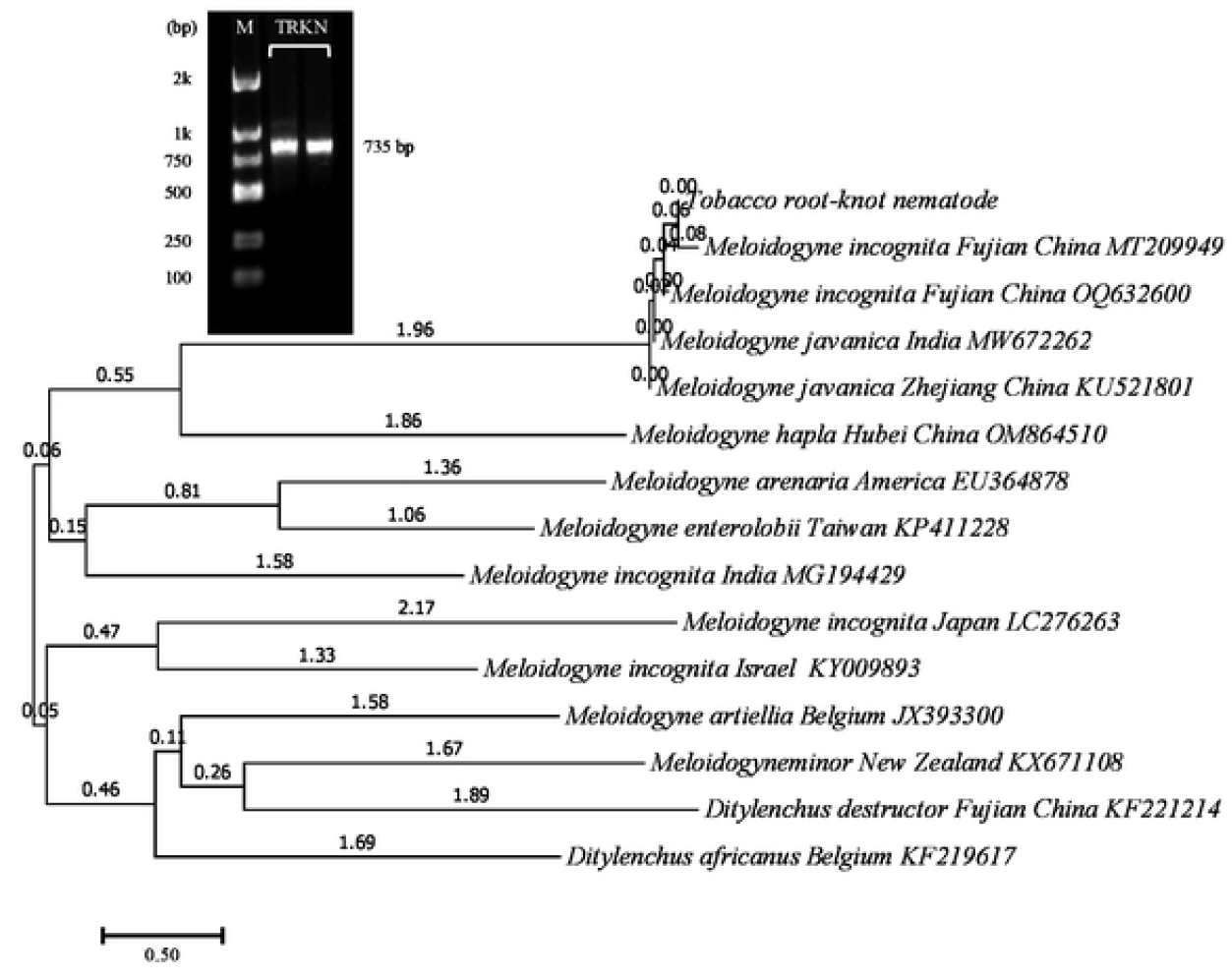
Phylogenetic relationships of the tobacco root knot nematode, *Meloidogyne incognita* and other selected *Meloidogyne* spp., as inferred from a 735 bp alignment of 28S rDNA sequences, with *Ditylenchus* genus as the outgroup.

Based on the ITS sequence of *M. incognita*, specific primers MiF and MiR were designed using the NCBI Primer-BLAST software. The forward primer exhibits a GC content of 38.1%, while the reverse primer has a GC content of 50%. A comparison of the primer sequences with the ITS sequences of other closely related species revealed that the forward primer had mismatches of 1, 6, 6, 10, 10, and 10 base pairs with *M. javanica* (MW672262), *M. minor* (KX671108), *M. artiellia* (JX393300), *M. hapla* (OM864510), *M. arenaria* (EU364878), and *M. enterolobii* (KZ411228), respectively. The reverse primer exhibited mismatches of 1, 3, 6, 10, 9, and 6 base pairs with the same species, respectively (Table 1).

**Table 1.**
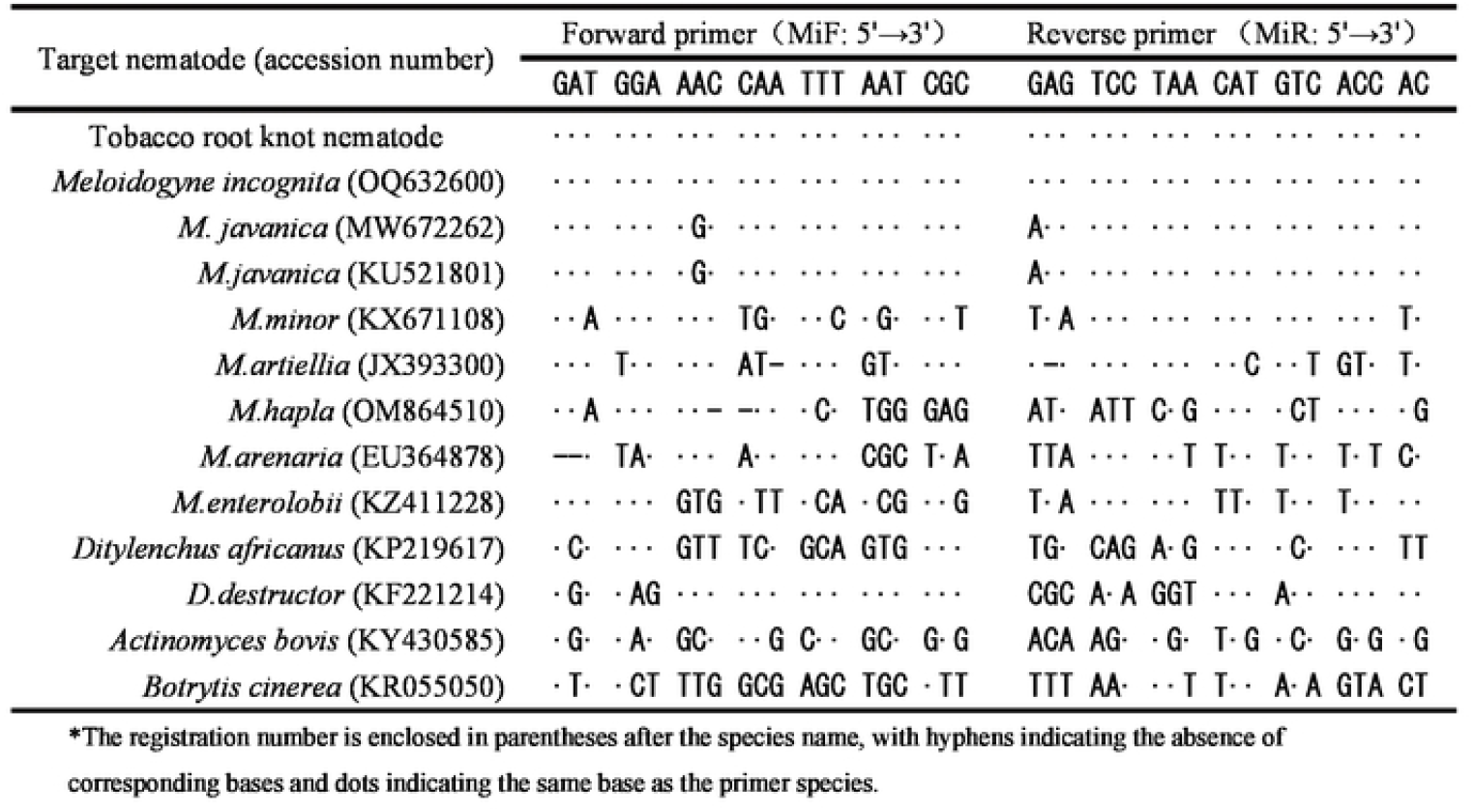
Sequence comparison between specific primers and related species ofTobaeco root knot nematode *(M incognito)*.

### 3.2 Ct value of different morphological nematodes

The egg masses extracted from the tobacco roots affected by nematodes at harvest time exhibited a milky white color and a smooth surface. Upon breaking open the egg masses for observation, three distinct morphological types of nematodes were identified: J1, which comprised eggs with clearly formed nematode bodies; J2, which were larvae that had hatched; and abnormal eggs, which were either hollow or contained bubbles (Figure 2). We extracted DNA from these different morphological types of nematodes and used them as templates for Ct value detection. The results indicated significant differences in Ct values among the various morphological types of nematodes. The Ct values for J1 extracted from the pre-planting soil and the affected roots at harvest were 19.6 to 19.9 (n = 5), respectively, showing no significant difference between the two. In contrast, the Ct values for J2 were lower than those for J1, recorded at 18.4 to 17.3 (n = 5), respectively. Notably, the Ct values detected from the abnormal egg DNA were substantially lower than those for J1 and J2, measuring 28.8 and 27.8, respectively, with a higher variance than that of J1 and J2. Furthermore, the egg masses extracted from the pre-planting soil and the affected roots at harvest were opened, and the eggs of each morphological type were counted. The results revealed that the proportion of abnormal eggs in the pre-planting soil was as high as 64.08%, whereas the proportion of abnormal eggs in the egg masses at harvest was 15.3% (Table 2).

**Table 2.** The Cl value *M incognila* by real-lime PCR at different developmental stages Life stage of *M*. From soil at pre-planting From root at harvest

| Life stage of *M. incognita* | From soil at pre-planting |  | From root at harvest |  |
| --- | --- | --- | --- | --- |
|  | Ct | individuals/egg mass | Ct | individuals/egg mass |
| Juvenile 1 | 19.6±2.1 a | 103.9±58.4 a | 19.9±0.2 a | 211.6±61.9 a |
| Juvenile 2 | 18.4±0.3 b | 15.0±8.0 b | 17.3±0.4 b | 231.9±67.6 b |
| Abnormal egg | 28.8±2.0 b | 211.6±61.9 c | 27.8±2.1 b | 78.0±18.0 c |
| Abnormal egg percentage/ egg mass | 64.08±16.7 % |  | 15.3±3.7 % |  |

\*The standard error in correction term is derived from standard error of the mean. Different lowercase letters after the data in the same column indicate significant differences among different treatments (n=5, $p < 0.05$ ).

**Figure 2.**
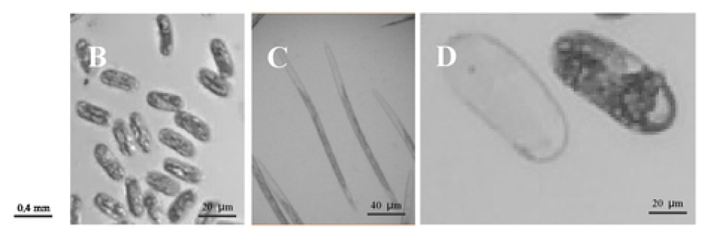
Morphology of egg mass on gall, eggs (both normal and abnormal), and Juvenile 2 (J2) stages of tobacco root knot nematodes *(M. incognila)* at various developmental stages. (A: the egg mass on galls at harvest, extracted from the diseased root of tobacco; B: JI, extracted from the galls on the root; C: J2, extracted from a suspension of nematodes; D: hollow and abnormal egg)

### 3.3 Quantitative Curve of M. incognita in Soil

We diluted the DNA solution of individual *M. incognita* as a template and detected the amplification of the designed specific primers MiF and MiR. The study revealed that the Ct values (y) increased logarithmically with the increase in the DNA dilution rate (x), with the regression equation defined as y = -0.9841x + 33.9857 and R^2^ = 0.9964. This indicates a highly significant correlation between Ct values and DNA dilution rate (P < 0.01). Furthermore, when the DNA was diluted to 10^5^ times, a linear relationship between the DNA dilution rate and Ct value was observed, with Ct values ranging from 19.8 to 32.9. However, when the dilution reached 10^7^ times, the Ct value was compromised, and no Ct values were detected in the negative control samples (Figure 3 A).

**Figure 3.**
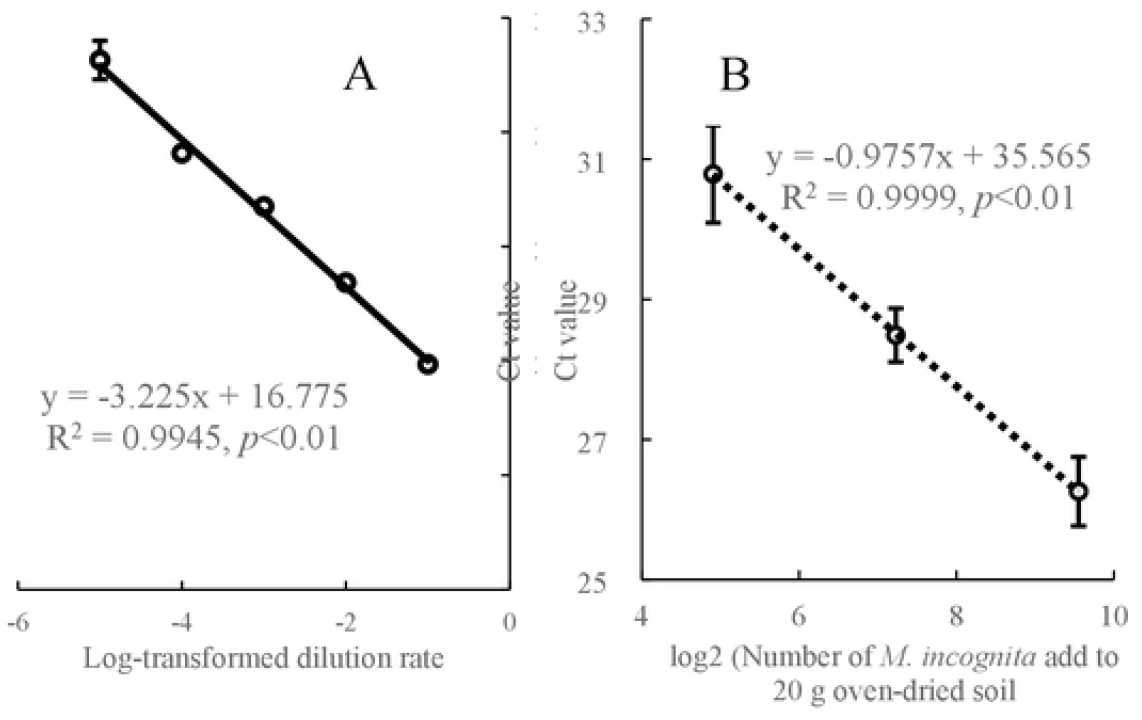
Effect of dilutions of a DNA ternplate extracted from nematodes infecting tobacco roots on the cycle threshold (Ct) values (A), and the relationship between the log-transfonned nun1bers of*M. incognito* added to 20 g of oven-dried soil and the Ct values (B; enor bars are standard deviation; n = 4).

In this study, based on the number of inoculated *M. incognita* in the soil and the Ct values detected by real-time PCR, a soil quantitative curve was constructed. A correlation analysis was performed between the Ct values (y) and the logarithmically transformed number of inoculated nematodes (x). The results revealed a significant negative correlation between the two, with the regression equation defined as y = -1.0859x + 32.025, R^2^ = 0.9866, P < 0.01 (Figure 3 B).

### 3.4 Degradation rate of DNA from M. incognita in soil

The initial inoculation density of dead *M. incognita* in the soil ranged from 2090 to 2275 individuals/20 g soil. By the third day, the DNA-based equivalent of nematode density decreased to 374±16 individuals/20 g soil, reflecting a degradation rate of 87.3%. On the seventh day,the DNA-based equivalent of nematode density further declined to 14±2 individuals / 20 g soil, indicating a degradation rate of 99.34% compared to the initial inoculation. By the fourteenth day, the DNA-based equivalent of nematode density in the soil had diminished to only 1.0±0.5 individuals/20 g soil, achieving a degradation rate of 99.97% relative to the initial inoculation (Figure 4).

**Fig. 4.**
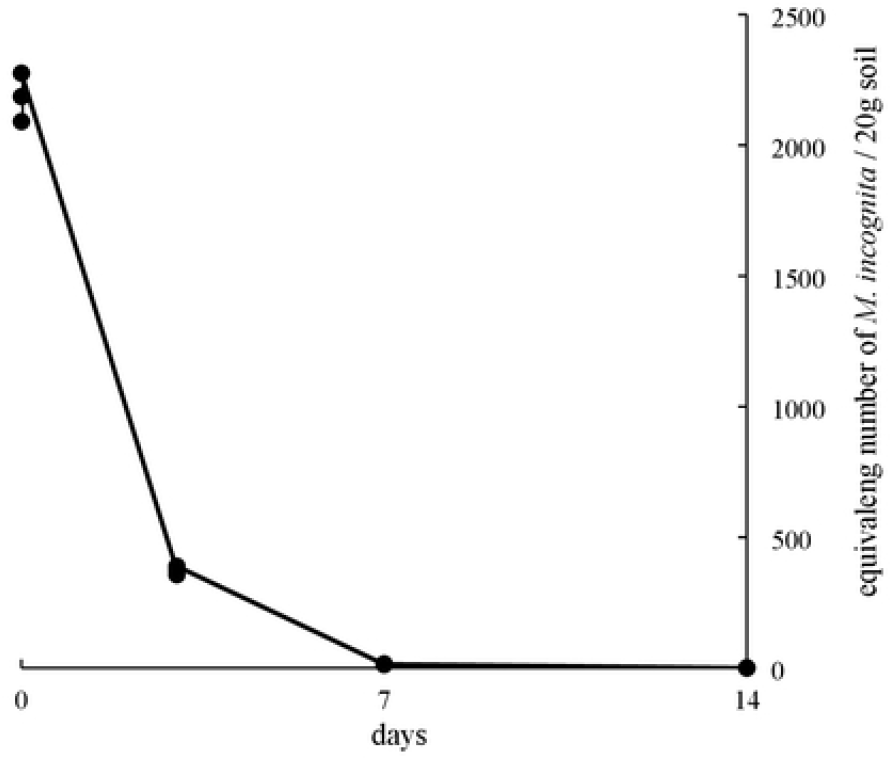
Decomposition rate of DNA from dead *M incognita* inoculated into 20 g of soil (n=3).

### 3.5 Effect of M. incognita density in the rhizosphere on the root-knot index

Following the tobacco harvest in October 2022 and October 2023, we conducted an investigation into the incidence of root diseases in 126 tobacco plants and assessed the density of *M. incognita* in the rhizosphere soil (Figure 5). The results indicated that at a Root-knot Index (RKI) level of 0, no galls or egg masses were detected in the tobacco root system; however, the density of *M. incognita* in the rhizosphere soil varied from 0 to 68 individuals/20 g soil. When the RKI reached level 1, the number of galls ranged from 2 to 9 galls/30 g root, with nematode densities in the soil ranging from 234 to 2321 individuals/20 g soil. At RKI level 2, the number of galls increased between 13and 23 galls/30 g root, with nematode density recorded at 2345-6764 individuals / 20 g soil. At RKI level 3, the count rose to 27 to 49 galls/30 g root, and the nematode density in the soil ranged from 4656 to 23434 individuals/20 g soil. At RKI level 4, the number of galls ranged from 53 to 73 galls/30g root, with a corresponding nematode density of 8233 to 35291 individuals / 20 g soil. Finally, at RKI level 5, the number of galls or swellings ranged form 71 to 86 galls/30 g root, with nematode density in the soil falling between 18345 and 49234 individuals/20 g soil. There were extremely significant differences between RKI level 0 and levels 1-5 (P < 0.01). Significant or extremely significant differences were found between RKI level 1 and levels 2-5 as well as between RKI levels 2 and 3-5, RKI levels 3 and 4-5, and RKI levels 4 and 5 (P < 0.05). Figure 5 showed that at RKI levels 1-3, as the number of galls increased, the nematode density in the soil also significantly increased, indicating a notable enhancement in significance. However, at RKI levels 4-5, the significance of this growth decreased, indicating that the variation in nematode density in the soil was reduced when the root-knot disease was severe.

**Fig. 5.**
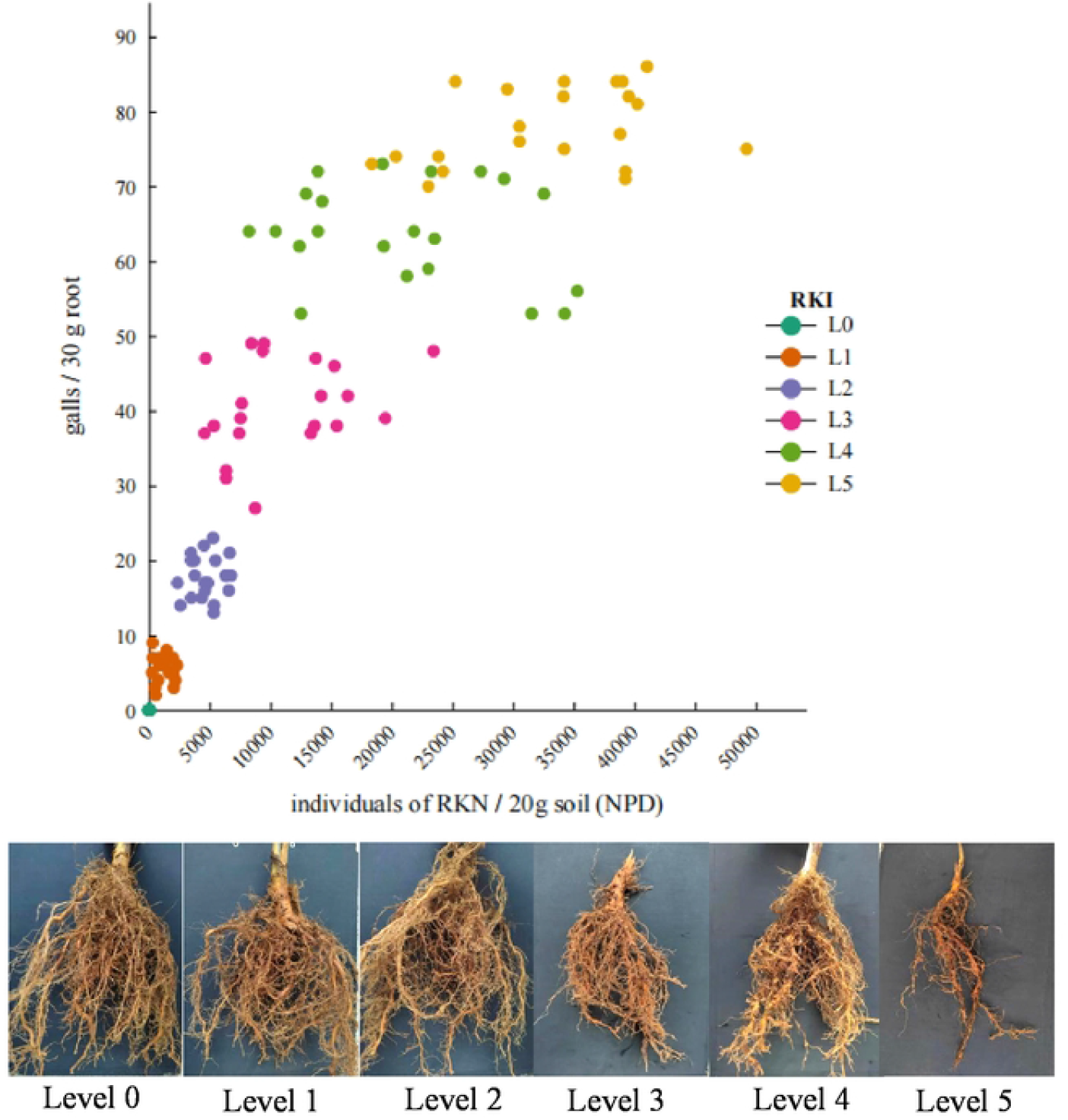
The relationship between the number of galls on 30 g root and the density of *M incognita* in the soil (RKI depends on gallson the tobacco root).

**Fig. 6.**
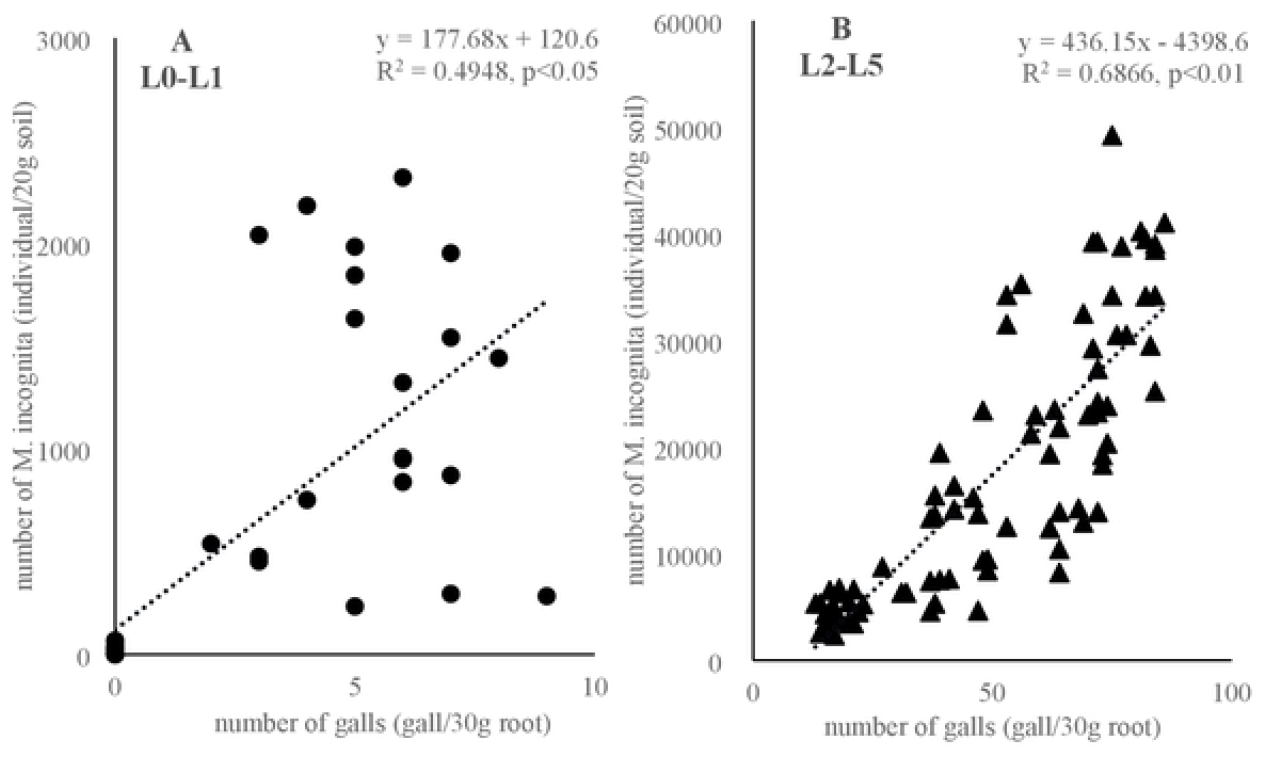
The relationship between the density of *M incognita* in the soil and the number of galls on root (low RKI: 0-1; High RKI: 2-5).

## 4. Discussion

This study successfully established a real-time PCR analysis method for tobacco root-knot nematodes (*M. incognita*) in the western region of Henan Province, demonstrating that the population density of *M. incognita* in the soil significantly impacts the occurrence of root-knot disease. Furthermore, the nematode samples collected from four tobacco-growing areas in this region exhibited complete consistency in their ITS region sequences, all identified as *M. incognita*. However, when comparing the sequences of different *Meloidogyne* species retrieved from the NCBI database with our designed specific primers MiF and MiR, 1-10 base mismatches were identified. According to Ri, M.U. et al. (2023), a forward primer with 10 base mismatches at the 3’ end significantly reduces amplification efficiency [32]. Moreover, if there are three or more base mismatches at the 5’ end, amplification will not occur. The *M. incognita* specific primers designed by Toyota et al. (2010) share identical sequences with *M. arenaria and M. javanica*, preventing differentiation between these species [21]. Similarly, the primers designed by Zhao et al. (2010) share the same sequence with *M. javanica* but differ by a single base from *M. arenaria*, resulting in an absence of observable PCR amplification products [33]. Additionally, Wu et al. (2024) reported that *M. arenaria* is the dominant population among Guizhou *M. incognita* [34]. Xu et al. (2023) indicated that in the primary tobacco-growing area of Kunming, Yunnan, *M. incognita* comprises 89.0% of the tobacco root-knot nematode population, while *M. arenaria* constitutes 41.3%, *M. javanica* 53.2%, and other *Meloidogyne* sp. 12.8% [35]. Jiao et al. (2014) reported the population distribution ratio of tobacco root-knot nematodes in Henan Province, revealing that *M. incognita* constituted at 55.83%, *M. arenaria* 23.33%, *M. hapla* 17.50%, and *M. javanica* 3.33% [36]. This finding further confirms that *M. incognita* is the dominant species in re region [37]. In this study, the *M. incognita* identified from tobacco samples collected in the Luoyang area of Henan Province was confirmed through ITS region sequence analysis, aligning with Jiao et al. (2014), who also identified *M. incognita* as the predominant tobacco root-knot nematode in Henan [36]. Concerning the closely related species *M. javanica*, our primers MiF and MiR exhibited one base mismatch each in their forward and reverse sequences. When we used the nematode DNA for detection using the MiF and MiR primers, the Ct value increased by 6, and the amplification efficiency decreased by 64% (data not shown), indicating a high specificity of these primers for *M. incognita*.

This study is the first to detect DNA in nematodes of varying morphologies within the egg mass, using the MiF and MiR primers for real-time PCR analysis of Ct values. The results indicated that the Ct values of abnormal eggs were significantly lower than those of normal eggs (J1), suggesting that these abnormal eggs are inactive and that their DNA has begun to degrade, rendering them incapable of hatching. This finding suggests that the hatching rate of *M. incognita* eggs in the soil during fallow periods, in the absence of a host plant, will significantly decrease. Additionally, this study examined the decomposition rate of DNA from deceased nematodes and found that seven days post-mortem, the amplification signal of their DNA was extremely low, nearly negligible. Consequently, the number of nematodes detected in the soil by real-time PCR could be interpreted as representing active nematodes or viable eggs. Furthermore, this indicates that even if *M. incognita* eggs hatch into J2, the rate of DNA abnormalities remains relatively high in the absence of a host plant. In conclusion, implementing fallow or crop rotation measures can effectively reduce the population density of *M. incognita* in the soil and significantly impact disease control. These results provide crucial scientific evidence for managing *M. incognita* and advocate for the adoption of fallow or crop rotation strategies in practical applications to mitigate nematode damage.

The results of this study on the relationship between RKI and NPD indicate a positive correlation between the number of *M. incognita* in the tobacco rhizosphere soil and the RKI levels of 1 to 3. This finding aligns with previous research reports, suggesting that during the early to mid-stages of the disease, the growth of the nematode population is significantly related to the severity of root-knot disease. In the initial stages of the disease (RKI 1-3), an increase in RKI levels corresponds with a significant rise in the number of nematodes within the rhizosphere soil. This indicates that following infection of the tobacco root system, nematodes could reproduce rapidly and establish a stable population. At this stage, the growth of the nematode population is primarily driven by the nutritional supply from the host root system, and the formation of root knots provides additional feeding sites for nematodes, thereby further promoting their reproduction [38]. However, as the disease progresses to higher levels (RKI 4-5), the nutritional capacity of the tobacco root system begins to decline, which restricts further nematode reproduction and results in a significant slowdown in the growth rate of the nematode population. Furthermore, the number of nematodes in the rhizosphere soil may have already reached the environmental carrying capacity, thus limiting the contribution of further increases in disease levels to the growth of the nematode population.

The findings of this study provide scientific evidence for the integrated management of tobacco root-knot nematode disease. In the early stages of the disease (RKI 1-3), the nematode population has not yet reached the environmental carrying capacity. At this stage, implementing control measures, such as the application of nematicides or the cultivation of resistant varieties, could effectively curb the increase in nematode numbers and mitigate the disease’s negative impact on tobacco yield []. Once the nematode population reaches a specific threshold, its growth rate significantly declines. Therefore, monitoring the nematode population in the rhizosphere soil during agricultural production can help determine the economic threshold for disease management, allowing for the rational use of pesticides, thus reducing production costs and minimizing environmental pollution. The integration of agricultural management practices, such as crop rotation and soil disinfection, along with biological control methods including the application of antagonistic microorganisms or entomopathogenic nematodes, can effectively control nematode population growth, reduce reliance on chemical pesticides, and enhance the sustainability of tobacco production.

Despite revealing the association between RKI and NPD, this study has several limitations. It did not account for the influence of environmental factors such as soil type and climatic conditions, on the dynamics of the nematode population. Future research should further investigate the specific roles of these factors. Additionally, genetic variations among *M. incognita* populations across different regions may impact their host infection capabilities and reproductive efficiency.

## 5. Acknowlegements

Funding for this study was generously provided by the Science and Technology Project Fund of the China Tobacco Corporation of Luoyang #2022410300270069, the Science and Technology Project Fund of the China Tobacco Corporation of Xuchang #2024411000240026, the Key Scientific and Technological Projects of the China Tobacco Corporation #110202201026 (LS-10), and the Henan Province Science and Technology Research Project #22010174. The isolates of the tobacco root-knot nematode and tobacco plant samples were collected in 2023 by Mr. Pu Miao from the China Tobacco Corporation of Luoyang.

## Notes

### Competing Interest Statement

The authors have declared no competing interest.

